# A Convenient Non-harm Cervical Spondylosis Intelligent Identity method based on Machine Learning

**DOI:** 10.1101/264663

**Authors:** Nana Wang, Xi Huang, Yi Rao, Jing Xiao, Jiahui Lu, Nian Wang, Li Cui

**Affiliations:** Institute of Computing Technology(ICT), Chinese Academy of Sciences(CAS), Beijing, China; University of Chinese Academy of Sciences, Beijing, China; Xiyuan Hospital, China Academy of Chinese Medical Sciences(CACMS), Beijing, China

## Abstract

Cervical spondylosis(CS), a most common orthopedic diseases, is mainly identified by the doctor’s judgment from the clinical symptoms and cervical change provided by expensive instruments in hospital. Owing to the development of the surface electromyography(sEMG) technique and artificial intelligence, we proposed a convenient non-harm CS intelligent identify method EasiCNCSII, including the sEMG data acquisition and the CS identification. For the convenience and efficiency of data acquisition with the limited testable muscles provided by the sEMG technology, we proposed a data acquisition method based on the relationship between muscle activity pattern, the tendons theory and CS etiology. It is easily performed in less than 20 minutes, even outside the hospital. Faced with the challenge of high-dimension and the weak availability, the 3-tier model EasiAI is developed to intelligently identify CS. The common features and new features are extracted from raw sEMG data in first tier. The EasiRF is proposed in second tier to further reduce the data dimension and improve the performance. With the limited and weakly available data, the gradient boosted regression tree is developed in third tier to effectively identify CS. The EasiAI achieve the best performance with 91.02% in accuracy, 97.14% in sensitivity, and 81.43% in specificity compared with 4 common machine learning classification model, validating the EasiCNCSII effectiveness.

---

Cervical spondylosis(CS) is a degenerative disorder common which affect up to two-thirds of the population in their lifetime^1–4^. The CS seriously affect people’s physical and mental health and quality of life and increase the burden on individuals and society. What’s more, it leads to the loss of human-related functions and is accompanied by depression, anxiety and other psychological damage. The main complaint of CS is neck pain which is reported by approximate 30-50% from the patients^5^. Meanwhile, the global point prevalence of neck pain was 4.9% and the neck pain ranked 4th highest in terms of disability as measured by YLDs, and 21st in terms of overall burden in the Global Burden of Disease 2010 Study^6^. The early detection of the CS is critical for burden lighten. As the earlier the disease is discovered, the easier it is to treat, the higher the cure rate is, and the less the patient spend. The CS is a chronic ‘wear and tear’ degenerative process of the cervical spine that initially is the vertebral bodies and intervertebral disks degeneration in the neck, and can develop into disk ruptures and herniation, osteophyte, compression of the spinal cord, or cervical spondylotic myelopathy(the most important complication of degenerative disease of the cervical spine)^7–9^. As cervical degeneration worsens, clinical manifestations become more obvious, and the difficulty and cost of treatment increases. In detail, for the patients with the vertebral bodies and intervertebral disks degeneration in the neck, non-operative treatment continues to play an important role in treatment^2^. For the patients with cervical spondylotic myelopathy, surgical treatment have been conventional means but may lead to significant problems including adjacent level^2, 10^.

Due to the complex of the pathogenesis and clinical symptoms of the CS, the identification of CS is a sophisticated and complicated work. A large number of auxiliary inspection methods are used to assist the identification. The following methods^11–16^ included in the domestic and foreign objective examination of the CS are clinical examinations, spinal angiography, vertebral artery angiography, X-ray, computed tomography(CT), and magnetic resonance imaging(MRI), etc. Most of them depend on expensive medical instruments in hospital to directly observe the physical changes in the spine and ancillary structures. However, the physical changes above can cause chronic dysfunction as well as pain^17^, leading to anomalous pattern of muscle activity^18–24^. And when the muscles are activated in the activity, the Motor Unit Action Potential Trains(MUAPTs) are generated by motor units, superimposed on the surface of the skin and form a non-stationary week signal which can be acquired by the sEMG device and generate electromyography. The relevant works^18–24^ demonstrated that there are differences in sEMG signals between population with cervical musculoskeletal disorders or neck pain and the healthy. Thus, it provides a chance that we can explore the relationship between sEMG signals and CS to identify CS. Benefit from the development of sensors technology, sEMG device become more portal and more cheaper, promoting the sEMG technology to become a competitive choice for the convenient CS identification. The sEMG have attracted a lot of attention in muscle function assessment^25, 26^, muscle activity assessment^20, 21, 23^, rehabilitation effect tracking^27^ and rehabilitation guidance^28^. Meanwhile, the development of machine learning has made great progress in medical^29–32^. With the convenient of sEMG technology and the development of artificial intelligence, the convenient, no-harm, intelligent CS identification method can be considered.

The data acquired by portal sEMG device present a huge challenge to the identification of CS. The high-dimensional sEMG data can cause dimensional disaster which decrease computational efficiency, increase memory storage requirements, and cause overfit. Faced with the high-dimensional data, feature extraction and feature selection, which are effective means of data preprocessing, have the advantages of improving model performance, increasing computational efficiency, decreasing memory storage requirements, giving model better readability and interpretability, and building better generalization model^33^. Besides, the data shows weak availability of faulty, redundant, insufficient, sparse distribution since the data acquisition is susceptible to external factors in non-lab environments by portal sEMG device. So a powerful machine learning model should be considered. Ensemble learning containing a number of weak learners can not only learn linear and complex nonlinear function but also boost weak learners which are slightly better than random guess to strong learners which can make very accurate predictions^34^. The gradient boosted regression tree (GBRT), one of powerful ensemble learning, has been successfully used in classification task^35–38^. And it is a competitive choice for the classification task on limited weakly available data.

In this work, we propose a new convenient, non-harm and intelligent method EasiC-NCSII to identify CS based on sEMG and machine learning as the Figure 1 shown. The method mainly consist of data acquisition and CS identification. For data acquisition, we proposed a convenient, time-saving data acquisition method. The subject only need to spend less than 20 minutes independently completing a set of simple movements according to instruction(see the Supplementary for the instruction of data acquisition), after connecting the portal sEMG device to the laptop and subject’s neck muscles. The relevant data is uploaded to the intelligent processing terminal while being collected by the sEMG device. For CS identification, the EasiAI model based on 3-tier architecture, was developed to identify CS. The EasiAI consist of feature extraction, feature selection and classification algorithm. For feature extraction, we extract 11 types of features from raw sEMG signal, of which 6 types are extracted in the common high dimensional time series feature extraction methods, such as time-domain method, of which 5 types are built inspired by the relevant knowledge (see the Supplementary for Feature extraction). Most of the features are proved to be significantly associated with the CS by Pearson(*p ≤* 0.05). For feature selection, EasiRF, a feature selection method, was developed to select the most relevant features and improve the performance of the CS identification. The easyRF is validated effective compared with traditional feature selection algorithms. For classification algorithm, a classification model based on GBRT is developed to identify CS. The EasiAI achieve the best performance with 91.02% in accuracy, 97.14% in sensitivity, and 81.43% in specificity compared with 4 kinds of machine learning classification model. The EasiCNCSII is validated effective.

**Figure 1:**
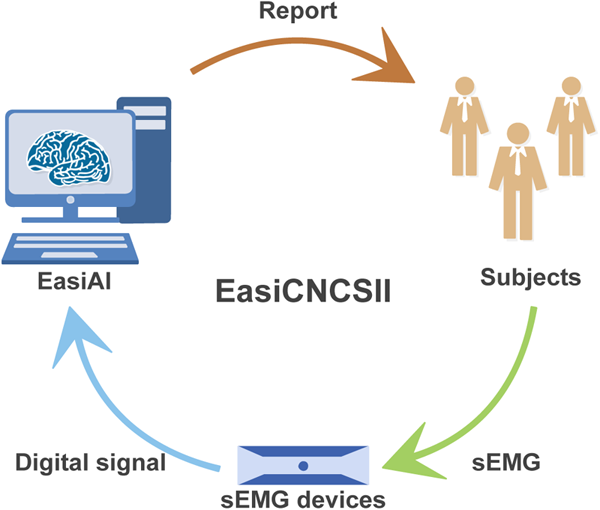
The CS identify based on sEMG and machine learning. Firstly, subjects perform a set of movements according to instructions. Secondly, the portable sEMG device acquire sEMG signals of the subject and send it to the smart terminal. Finally, the EasiAI, an intelligent CS identification model, predict the state of subject using the sEMG signals and return the report which shows whether the subject suffer from the CS.

## Result

### Data acquisition

The 537 samples are obtained, of which 327 are from patient and 210 are from the healthy free from the CS, since the 179 subjects are recruited in this work, of which 109 are patient and 70 are heathy, and a set of movements are repetitively completed three times by each subject.

The data acquisition method involve 6 muscles and 7 movements. The muscles above consist of the the left sternocleidomastoid(*M*_1_), left upper trapezius(*M*_2_), left cervical erector spinae (*M*_3_), right cervical erector spinae(*M*_4_), the right upper trapezius(*M*_5_), the right sternocleidomastoid(*M*_6_) (see Supplementary Figure *S*_1_ for the location of surface electrodes). The movements above include bow(*A*_1_), head backwards(*A*_2_), left flexion(*A*_3_), right flexion(*A*_4_), left rotation(*A*_5_), right rotation(*A*_6_), hands up(*A*_7_)(see Supplementary Figure *S*_2_ for the instruction of the movements). Each movement is held for 20 seconds and repeated 3 times with 5 seconds rest between each repetition.

After connecting the sEMG device to the subject’s neck muscles above as well as the laptop, the data acquisition begins. The subject performed the 7 movements according to instruction(see Supplementary for the instruction of data acquisition). After data acquisition, we obtain 7 multiple high-dimensional time series data from a sample, each of which is generated from the movement *A*_*i*_ and represented as *S*_*i*_. The *S*_*i*_ is expressed as Equation 1. The *S*_*i,j*_ denotes the sEMG signals collected from the muscles *M*_*j*_ activated by movement *A*_*i*_, expressed as Equation 2.

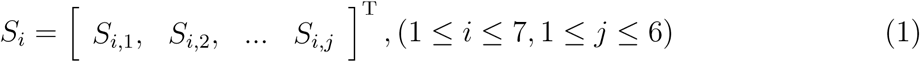

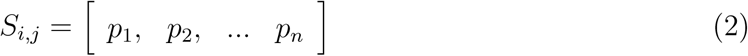

Here, *S*_*i,j*_ converted from the electrical signal are high dimensional time series data. The *p*_*n*_is a value of *S*_*i,j*_.

### Feature extraction

The 5 common methods: time-domain, frequency-domain, time-frequency, parametric model and nonlinear feature analysis are used to extracted features from the sEMG signal which are high-dimensional time series data(details on the calculation are shown in Supplementary Table *S*_2_). Besides, the method to build features using the disease-related knowledge are also considered. With the 5 common methods above, we extracted 63 features from *S*_*i,j*_, of which 11 are extracted in methods of time-domain^39, 40^, of which 14 are computed in methods of frequency-domain^39, 40^, of which 23 are computed in methods of time-frequency based on wavelet transform and wavelet packet transform^41, 42^, 14 of which are extracted in AR parametric model ^43^, of which 1 nonlinear entropy feature^39, 40, 44^ are extracted. Among the 63 features above, the 5 features including the root mean square (RMS), median frequency(MF), mean power frequency (MPF), the average electromyogram (AEMG), and the integrated electromyogram (IEMG) are common features in clinical research. Besides, considering the relative knowledge of the CS, the 45 features are extracted from *S*_*i*_, of which 2 called as cervical flexion-relaxation ratio(FRR)^45, 46^ is commonly used in clinical research and can only be extracted from the data of *S*_1_, and of which 43 are new features (see Supplementary for Feature extraction). The 63 features from *S*_*i,j*_ and 45 features from *S*_*i*_ are divided into 11 types: TF, FF, WL, WLP, AR, EY, FRR, DU, ACI, UN, SYM (details are shown in Supplementary Table *S*_1_-*S*_2_), facilitating statistical analysis.

The 423(45 + 6 *×* 63) features are extracted from *A*_1_ and 421(43 + 6 *×* 63) features^1^ are respectively extracted from the other 6 movements. Thus, 2949(423 + 6 *×* 421) features are extracted from the raw sEMG signal generated by all the movements above^2^ (the 2949 features’s distribution are shown in Supplementary Figure *S*_4_). The Pearson correlation indicated that 1789 features are significantly associated with the CS (*p ≤* 0.05, the significant features’s distribution are shown in Supplementary Figure *S*_4_).

### Data preparation

The data set consist of 537 samples and the number of features in each sample is 2949. We split the data into training samples(training set), validation samples(validation set), and test samples(test set) according to 16: 4: 5. Meanwhile, it is ensured that samples belonging to the same subject are only distributed in one of the three sets above.

### Feature Selection

We developed a feature selection algorithm EasiRF based on random forest(RF) to select the most relevant features and improve the performance of the CS identification. The EasiRF divide the 537 samples into 7 different data sets, each of which is represented as *D*_*i*_(0 *≤ i ≤* 7). The RF(with different number of trees) is iteratively used to select the top 25 most important features from *D*_*i*_. And put the 25 selected features of each iteration into the feature set *S*_*i*_ until the feature number of the *S*_*i*_ is not growing. As shown in Supplementary Figure *S*_5_, the selected feature number tends to be stable on each data set *D*_*i*_ when the iteration number reaches 40. The final feature set including 282 features are generated after merging all the *S*_*i*_ above(the feature type distribution of 282 features are shown in Supplementary Figure *S*_6_).

In order to validate the effectiveness of the EasiRF, the Fisher Score(FS)^47^, Conditional Infomax Feature Extraction(CIFE)^48^, Multi-Cluster Feature Selection(MCFS)^49^, f-score are respectively selected from 4 kinds of traditional feature selection algorithms^33^ as well as EasiRF. With the metrics of accuracy, sensitivity, specificity, FNR and FPR (see Supplementary for The metric of model performance), we compare the performance of the GBRT on test set with the feature selection algorithms above, using five-folds cross-validation. The large the accuracy, sensitivity and specificity are, the smaller the FNR and FPR are, the better performance of GBRT is, the more efficient the feature selection algorithm is. As shown in table 1, the mean accuracy of the model is 86.54% without feature selection. In spite of the slight drop in mean accuracy with the MCFS and CIFE, the mean accuracy of other algorithms is over 87.10%. What’s more, the mean accuracy with the EasiRF is the highest with the value of 91.02%. The mean sensitivity of the model is 92.51% without feature selection. All the mean sensitivity of the model are over 92.51% with feature selection algorithms. And the mean sensitivity with the EasiRF is the highest with the value of 97.14%. The mean specificity is 77.14% without feature selection. However, the mean specificities with feature selection algorithms are less than 77.14%, except the EasiRF with the highest value of 81.43%. The mean FNR of the model is 7.49% without feature selection. All the mean FNR with the feature selection algorithms are less than 7.49%. The mean FNR with the EasiRF is the smallest with the value of 2.86%. The mean FPR of the model is 22.86% without feature selection. And the mean FPR with feature selection algorithms are larger than 22.86%. The FPR with EasiRF are the smallest with 18.57%. Compared with the feature selection algorithms above, the EasiRF perform best and are validated most effective. The comparison of more feature selection algorithms are shown in Supplementary Tables *S*_3_.

**Table 1:** The comparison of performance with different feature selection algorithms

|  | Accuracy | Sensitivity | Specificity | FNR | FPR |
| --- | --- | --- | --- | --- | --- |
| Non | 86.54% | 92.51% | 77.14% | 7.49% | 22.86% |
| FS | 87.10% | 95.28% | 74.29% | 4.72% | 25.71% |
| CIFE | 86.00% | 92.60% | 75.71% | 7.40% | 24.29% |
| MCFS | 83.16% | 92.60% | 68.57% | 7.45% | 31.43% |
| F-score | 87.10% | 95.28% | 74.29% | 4.72% | 25.71% |
| EasiRF | 91.02% | 97.14% | 81.43% | 2.86% | 18.57% |

### the train and test of the EasiAI

We train the EasiAI on training set and validate on validation set. With larger AUC (the area under the sensitivity and specificity curve) values indicating higher classification accuracy, sensitivity and specificity across a range of threshold choices^29, 50^, the AUC is used to illustrate the performance of the classifier. The performance of the classifier is mainly affected by model parameters, especially the number of weak classifiers. Thus we first assessed that the number of trees(weak classifiers) included in the EasiAI were enough to obtain the highest AUC on validation set. As shown in Supplementary Figure *S*_7_, the higher the number of trees is, the higher the AUC on the validation set is. Furthermore, the AUC on the validation set achieved highest when the tree number is 535. The other parameters of the EasiAI model are shown in Supplementary Table *S*_4_. The final model is generated by training EasiAI with the parameters in Supplementary Table *S*_4_ on the dataset consisting of the training set and validation set.

To validate the performance of the final EasiAI, we tested the model on test set, using five-fold cross-validation. The metrics are accuracy, sensitivity, specificity, FNR, FPR as well as AUC. The large accuracy, sensitivity, specificity and AUC are, the smaller FNR and FPR are, the better the model classification performance is. The AUC, the mean accuracy, the mean sensitivity, the mean specificity respectively are 0.95, 91.02%, 97.14% and 81.43%.

The mean FNR, the rate of missed diagnosis, is 2.86%. The mean FPR, the misdiagnosis rates, is 18.57%. Overall, the accuracy of model is higher than 90%, the missed diagnosis rate of our model is less than 3%, and is validated effective.

### Comparing with support vector machines(SVM), Logistic regression(LR), NativeBayes(NB), random forests(RF)

The identification of CS is a task of classification. Faced with limited data, we chose the following four models that are often used, which include support vector machines(SVM), Logistic regression(LR), Native Bayes(NB), and random forests(RF). We validate the effectiveness of the EasiAI by comparing the EasiAI with SVM, LR, NB and RF, on test set^3^ in the same classification task, using five-fold cross-validation. As shown in table 2, the highest mean accuracy is 91.02% achieved by the EasiAI, and the lowest mean accuracy is 82.10%. The highest mean sensitivity is 97.14% achieved by the EasiAI, and the lowest mean sensitivity is 88.92%. The highest mean specificity is 82.86% and the lowest mean specificity is 70%. The mean specificity of the EasiAI is 81.43%, just smaller than the highest. The highest mean FNR(false negative rate) is 11.08% and the smallest value is 2.86% achieved by the EasiAI. The highest mean FPR(false positive rate) is 30% and the lowest value is 17.14%. The mean FPR of the EasiAI is 18.57%, just larger than the smallest one. Besides, the curves of sensitivities and specificities of models above are shown in Figure 2. Most of the sensitivity and specificity point falls below the red curve of EasiAI, especially between point a and b. The EasiAI achieve a best performance compared with RF, SVM, LR and NB with the metrics above.

**Table 2:** The performance of machine learning on test set

|  | Accuracy | Sensitivity | Specificity | FNR | FPR |
| --- | --- | --- | --- | --- | --- |
| EasiAI | 91.02% | 97.14% | 81.43% | 2.86% | 18.57% |
| RF | 87.65% | 93.42% | 78.57% | 6.60% | 21.43% |
| SVM | 86.00% | 89.78% | 80.00% | 10.22% | 20.00% |
| LR | 86.57% | 88.92% | 82.86% | 11.08% | 17.14% |
| NB | 82.10% | 89.83% | 70.00% | 10.17% | 30.00% |

**Figure 2:**
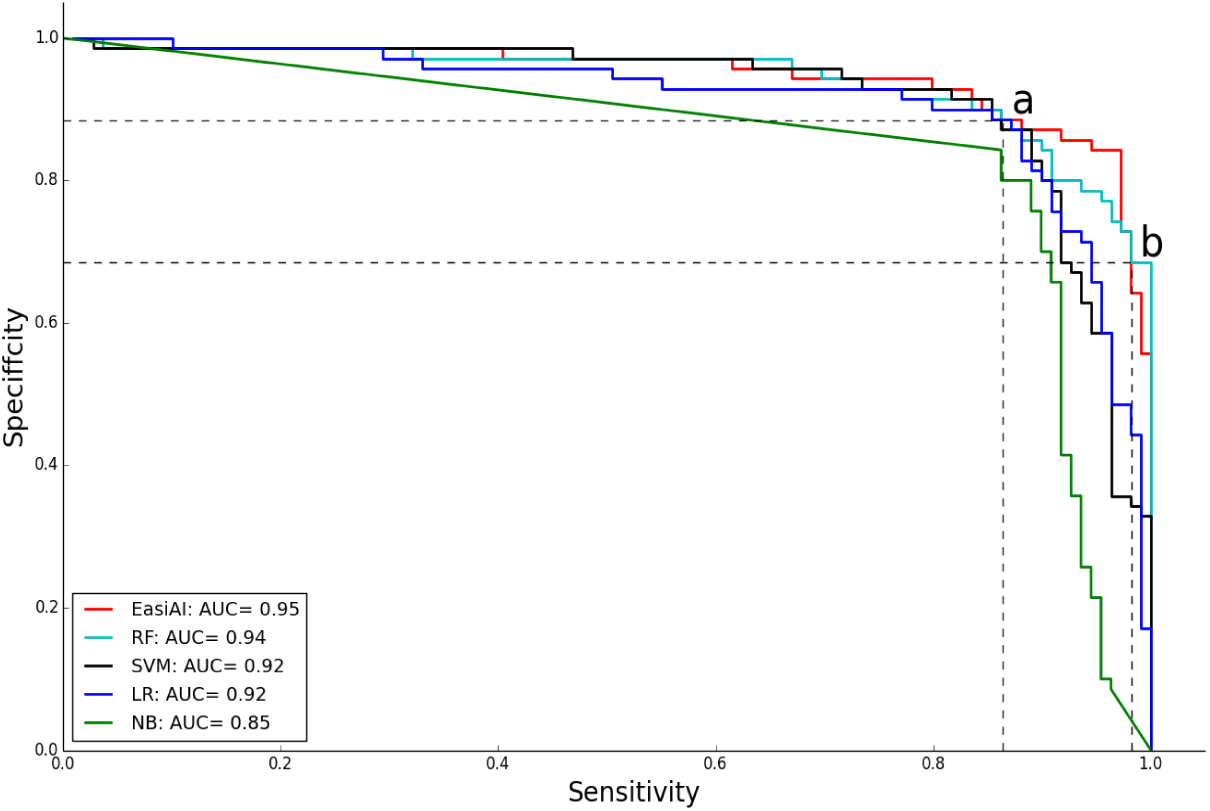
The performances of the EasiAI, RF, SVM, LR, and NB. The x-axis is the sensitivity. The y-axis is the specificity. Different color denotes different machine learning model. The curves of different colors consist of sensitivity and specificity points of different models. The closer the sensitivity and specificity points are to the upper right corner, the large the sensitivity and specificity are, the better the model performs. All the sensitivity and specificity points of the NB are the farthest from the upper right corner and fall below the curves of other models. The two curves of the LR and the SVM are intertwined whose most of sensitivity and specificity points fall below the curve of the RF. The two curves of RF and EasiAI are intertwined and most of the RF’s sensitivity and specificity points fall below the ones of EasiAI, especially between point a and b. The sensitivity and specificity points of the EasiAI are closest to the upper right corner, and have the highest AUC value 0.95.

### Analysis of learned knowledge about CS

The EasiAI achieved high accuracy in identifying CS, so we believe that the features in final model play an important role in the classification. And the informative knowledge about the CS can be learned from the feature distribution in muscles and movements. In order to further analyze the feature distribution above, the number^4^ distribution and importance^5^ distribution of the features in the muscles and movements are plotted as units of the heat map in Figure 3. The darker the color of the unit is in Figure 3, the more the features on the unit are in Figure 3 (a), and the more important the features on the unit are in Figure 3 (b). As shown in Figure 3, the features on the units formed by the *M*_3_(left CE) and *A*_1_(bow), *M*_4_(right CE) and *A*_1_(bow), *M*_2_(left UT) and *A*_7_(hands up), *M*_5_(right UT) and *A*_7_(hands up) are the most in number and most important for the CS identification performance. It is concluded that the CE activated by the movement of *A*_1_(bow head) and the UT activated by the movement of *A*_7_(hands up) may be more closely related to the CS.

**Figure 3:**
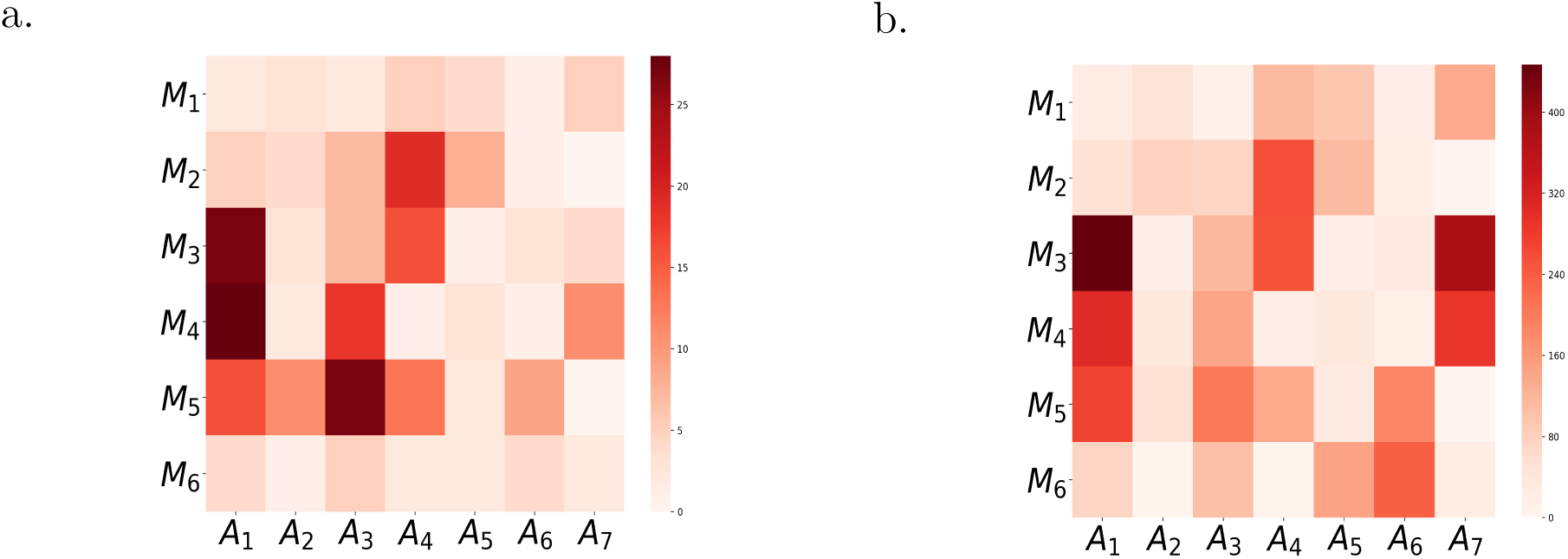
The heat map. The *M*_1_, *M*_2_, *M*_3_, *M*_4_, *M*_5_, *M*_6_ respectively denotes left SCM, left UT, left CE, right CE, right UT and right SCM. The *A*_1_, *A*_2_, *A*_3_, *A*_4_, *A*_5_, *A*_6_ and *A*_7_ respectively denotes low head, head backwards, left flexion, right flexion, left rotation and right rotation, hands up. (a), The number distribution of features in muscles and movements. A square represent the number of feature on *M*_*i*_ activated by *A*_*i*_. The darker the square’s color is, the more the features on the square are. (b), The importance distribution of features in muscles and movements. A square represent the importance of feature on *M*_*i*_ activated by *A*_*i*_. The darker the square’s color is, the more important the features on the square are.

In order to further understand feature distribution, the feature number distribution in muscles(A-distribution), the importance distribution in muscles(B-distribution), the feature number distribution in movements(C-distribution), and the features importance distribution in movements(D-distribution), are plotted as Figure 4. As shown in Figure 4 (a), the features on UT (*M*_2_ and *M*_5_) and CE (*M*_3_ and *M*_4_) take up 85.47% in the A-distribution and 74.60% in the B-distribution. As shown in Figure 4 (b), the features on the *A*_1_(bow) rank first in both C-distributions and D-distribution. The features on *A*_7_(hands up) rank second in D-distribution. It is concluded that the CE play an most important role in identifying CS followed by UT. And the movements of *A*_1_(bow head) also play an most importance role in identifying CS, followed by the movement of *A*_7_(hands up).

**Figure 4:**
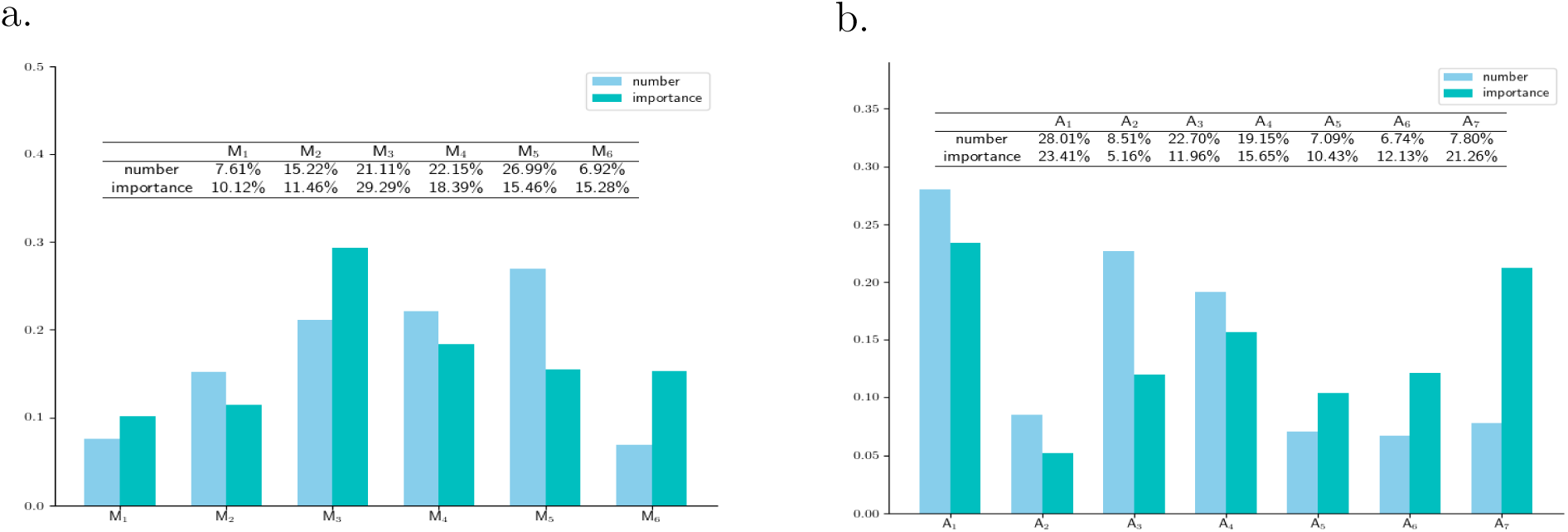
The feature distribution. (a), the feature distribution on the muscles. The *M*_1_, *M*_2_, *M*_3_, *M*_4_, *M*_5_, *M*_6_ respectively denotes left SCM, left UT, left CE, right CE, right UT and right SCM. The light bars show the distribution of feature numbers(A-distribution). The right UT rank first, followed by right CE, left CE, left UT, left SCM, and right SCM. The right UT, right CE and left CE take up 70.25%. The dark bars show the distribution of feature importance(B-distribution). The left CE rank first, followed by right CE, right UT, right SCM, left UT and left SCM. The left CE, right CE and right UT take up 63.14%. (b), the feature distribution on the movements. The *A*_1_, *A*_2_, *A*_3_, *A*_4_, *A*_5_, *A*_6_ and *A*_7_ respectively denotes low head, head backwards, left flexion, right flexion, left rotation, right rotation and hands up. The light bars show the distribution of numbers(C-distribution). The movement of head backwards rank first, followed by left flexion, right flexion, low head, hand up, left rotation, right rotation. The dark bars show the distribution of feature importance(D-distribution). The movement of the bow head rank first, followed by hands up, right flexion, right rotation, left flexion, left rotation, head backwards.

The conclusion can be explained by that the more frequently muscle are activated, the more the muscle(movement) is related to CS. Most of subjects are sedentary population who frequently bow head and use the hands and arms to work. The movement of bowing head activate the CE, and the movement of the hands and arms activate the UT. The excessive use of the CE and UT cause muscle strain, cause dysfunction of the spine stabilizing system, and accelerate cervical degeneration and lead to CS. It may provide a suggestion for sedentary population to prevent CS by avoiding excessive use of the CE and UT, strengthen the UT and CE protection.

### Discussion

The CS identification methods mainly include clinical symptoms examination and imaging examination since the diagnosis of CS is determined by clinical symptoms and imaging information^16, 51^. Currently, the clinical symptoms examination are performed by experts or doctors in the form of the inquiry. The images examination^6^ mainly depend on observation of the physical changes in spine and its subsidiary structure by imaging instruments to identify CS. The EasiCNCSII depend on detecting abnormal sEMG signal associated with muscle activity to identify CS. We compare the practicality of the EasiC-NCSII with the inquiry, imaging method as shown in table 3. The inquiry method is easiest and fastest. However, the method is suitable for population with obvious symptoms, for example severe pain, since information is mainly determined by suffer’s subjective feelings and judgment. And it needs the help of the doctor and auxiliary equipment to accurately identify CS. The images examination is the essential CS examination currently. However, its cost is relatively high and it is time-consuming considering time spent to go to the hospital and wait for the results which requires the intervention of doctors or experts. What’s more, it can not be used frequently with concern on health since the frequent use of imaging instruments can put a strain on the health, for example the radiation. Compared with inquiry and imagines examination, the EasiCNCSII is an best choice to identify CS outside the clinic with the advantage of easy use, low cost, no-harm. Due to the intelligent algorithm, users can get results quickly after examination. What’s more, combined with mobile application and wearable sEMG acquisition technology, the EasiCNCSII potentially provide low-cost convenient universal access to indispensable care outside the clinic, and even promote the development of telemedicine, especially in areas short of medical resources.

To our best knowledge, previous research on CS identification based on the sEMG and machine learning are few, so there is a lot room for improvement. Traditional classification or regression algorithms can achieve good performance when fed with a wealthy of high quality data. However, the amount of data is limited and the data dimension is high. Besides, data acquisition is vulnerable to the environment so that there are some poor quality data collected by portal sEMG. The EasiAI can handle these influences by boosting multiple weak learners to reach higher prediction accuracies. Compared with the RF, SVM, LR and NB, the EasiAI can achieve best performance, and can identify complex crowd with low missed diagnosis rate. It is hardly to fully understand the relation between CS and the sEMG signal from the activity of the deep and shallow muscles with the limited data, but the data-driven machining learning can achieve better performance with more accumulated data, and can accelerate our understanding of the principles behind the CS identification to assist diagnosis and guide treatment. We are looking forward to more convenient and intelligent applications in CS studies

## Methods

### Cohort description

The study population are made up of the outpatient from Beijing Xiyuan Hospital Physiotherapy Department and the healthy free from the CS from Institute of Computing Technology(ICT) and the hospital above, which were recruited as volunteer by email and advertising. A total of 179 subjects participated in the research, of which 109 were patient and 70 were heathy. The former are outpatients that have received a clinical diagnosis of the CS, which was in accordance with 2012 ICD-9-CM Diagnosis Code 721(721.0 Cervical spondylosis without myelopathy, 721.1 Cervical spondylosis with myelopathy) and the diagnostic criteria of diagnosis and treatment for CS issued by China Rehabilitation Medicine association. The latter are free from the CS diagnosed by the rich experienced clinicians. The exclusion criteria includes cervical vertebral trauma, cervical vertebral surgery, congenital spinal deformity, syringomyelia, amyotrophic lateral sclerosis, spinal cord tumor, spinal cord injury, adhesive arachnoiditis, the fascitis, cervical injury, tumor or infection, pregnant, breast-feeding, menstruation. All the subjects were given a clear explanation of tests approved the research ethics board of Beijing Xiyuan Hospital before taking part in this research.

### The selection of muscles and movements

In order to identify CS, the selected muscles should meet the following requirements. Firstly, the selected muscles are associated with the CS. It is known that CS belong to tendons injury in traditional Chinese medicine. It is supported by the tendons injury theory in traditional Chinese medicine that cervical soft tissue abnormalities cause CS^52^. The tender points are the indication of cervical soft tissue abnormalities^53^. The tender points of CS mostly focus on the cervical paravertebral muscle, trapezius and sternocleidomastoid muscle^54^. Thus, the cervical paravertebral muscle, trapezius and sternocleidomastoid muscle are competitive choice. Secondly, the selected muscles can be activated by the activities, producing sEMG signal. The CE and the SCM participate in most of the neck basic function activities above. The trapezius are activated by the scapula activities which mainly includes shoulder and hand movements, especially the upper trapezius(UT). Thus, the CE, SCM and UT are suitable choice. Finally, the sEMG signal generated by the selected muscles can be easily acquired with the minimal interference. The guidelines for electrode placement^55–58^ provide the mature surface electrodes location for CE, UT and SCM. Thus, we select the spinae (CE), the sternocleidomastoid (SCM) and upper trapezius (UT) as collected muscles. The electrode locations of the UT, CE, SCM, and reference ground are shown in Supplementary Figure *S*_1_.

In order to identify CS, the selected movements should meet the following requirement. The movement can maximally activate muscles to produce the most obvious sEMG signal which is easily collected by sEMG device and have the most significant difference between CS suffer and the healthy free from the CS. Since it is hardly to fully understand the principles between sEMG signals and the activity of the deep and shallow muscles to our best knowledge, so all the movement activating the muscles above need to be considered. The function activities, which CE, SCM and UT are involved in, are mainly rotation, lateral flexion, bow, head backwards and scapula activities. So we select 7 representative movements from the functional activities above which consist of the following movements: bow, head backwards, left flexion, right flexion, left rotation, right rotation and hands up. The movements are shown in Supplementary Figure *S*_2_.

### The EasiCNCSII

In this paper, we proposed a method EasiCNCSII to identify CS. As shown Figure 5, the method include two parts: data acquisition and the EasiAI model. For data acquisition, the subjects are connected sEMG device with the SCM, UT and CE and complete 7 movements according to the simple instructions. The analog signals from the subjects are converted into the digital signals, which are high dimensional time series data and sent to the EasiAI, by the sEMG device. The EasiAI is a three layer data processing model: feature extraction, feature selection and classification algorithm. Feature extraction algorithm extract 2949 features from the digital signals of a subject in the methods of time-domain, frequency-domain, time-frequency-domain, etc. Feature selection algorithm is developed to selected the most relevant features by iterative RF. The 282 features are selected from the 2949 features above. The gradient boosted regression tree is developed to identify CS, achieving the good performance on limited data set with relative small computing overhead. With the input data consisting of 282 features above, the report, which show whether the subject suffer from the CS, is generated and returned to the subject. The lightweight algorithms can be integrated into the user end, and quickly return report to subjects without concerns on privacy.

**Figure 5:**
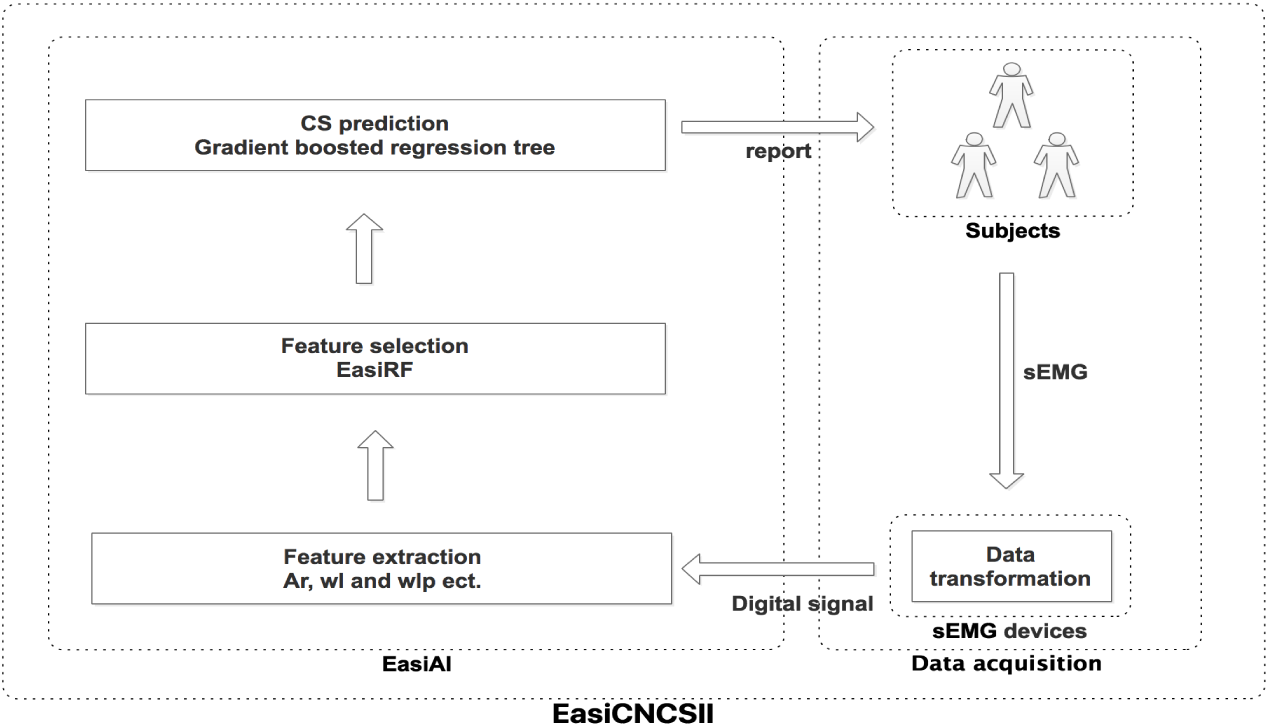
The EasiCNCSII method. The left side of figure is the CS identify model EasiAI consisting of feature extraction, feature selection, classification model. The right side of figure is data acquisition. The data collected by sEMG is automatically transmitted to the intelligent terminal equipped with EasiAI. The report generated by the EasiAI is sent to the subjects.

### The EasiRF algorithm

The EasiRF based on the RF is a stochastic model since the samples are randomly selected to generate the tree and the features are randomly selected to be splitting rule. Firstly, we generate 7 different data sets and iteratively use random forest to train on each data set. Secondly, set the tree number of RF model different, select the top 25 most important features and merge the features of each iteration until the feature number of the merged set is not growing in each iteration. Finally, the final feature set is generated by merging 7 feature sets from 7 data sets above. As shown in algorithm 1, the feature selection algorithm EasiRF based on RF is developed to get the most relative features.

### Experimental set-up

The python (version 2.7.13) and matlab (version 2016r) were used to implement features extraction from sEMG data. The python (version 3.6.0) and xgboost (version 0.6) were used to implement the EasiAI model. The RF, NB, LR and SVM model are implemented by the scikit-learn(0.19.1) and the python (version 3.6.0). The laptop and sEMG device which used to collect the sEMG signal are provided by Wireless Sensor Network Lab(this research does not develop the hardware). The EasiAI is deployed in the laptop with the CPU of i7, the memory of 8GB and operating system of 64-bit.

**Table 3:**
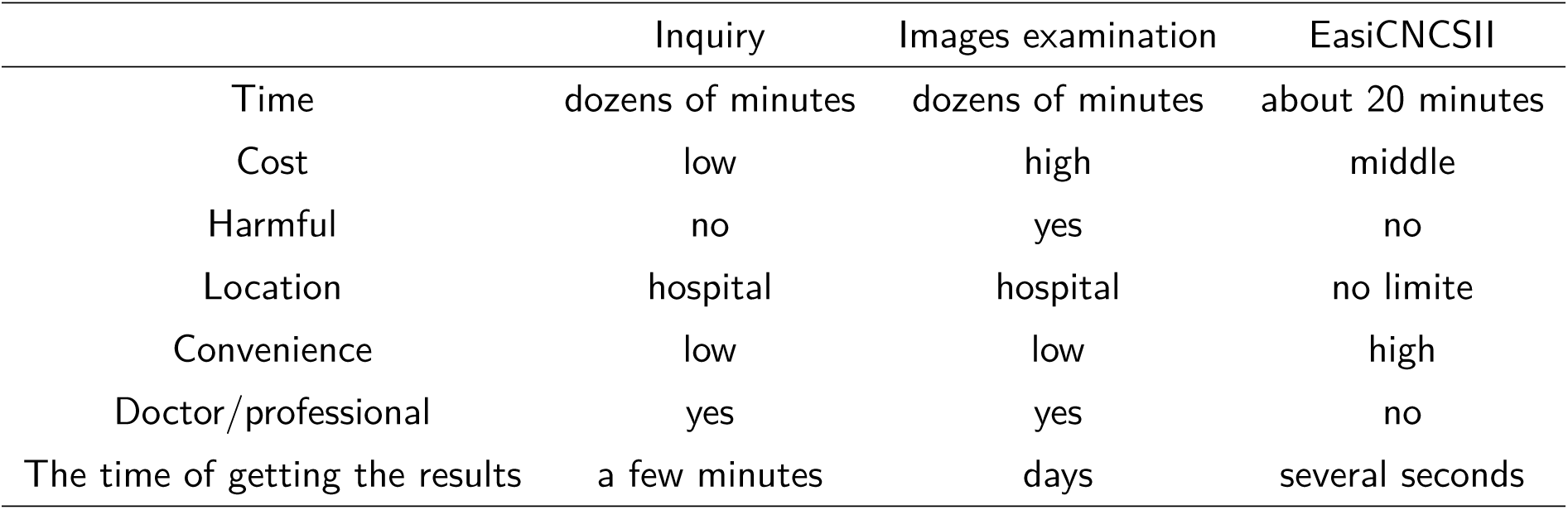
The comparison of the Inquiry, Images examination, EasiCNCSII

#### Algorithm 1 The EasiRF algorithm

~~~
**Input:** the data set *S*
**Output:** the set of features *FeaSet*
1:*k*=7, *FeaSet* = Ø
2:Divide the *S* into *S*_1_, *S*_2_, *S*_3_,…, and *S*_7_
3:**for** *j* = 1 to 7 **do**
4:  *DS* = *S* - *S*_*j*_
5:  *FeaSet*_*j*_ = Ø
6:  *Num* = 0
7:  **while** *| FeaSet*_*j*_ | is not growing **do**
8:    *FeaSet*_*t*_ = Ø
9:    Set the number of the RF with (50 + *Num ∗* 5).
10:   Run the RF with *DS*, output the features and its importances.
      Put the 25 the most important features into *FeaSet*_*t*_
11:   *FeaSet*_*j*_ = *FeaSet*_*j*_ *∪ FeaSet*_*t*_
12:   *Num* = *Num* + 1
13: **end while**
14: *FeaSet* = *FeaSet ∪ FeaSet*_*j*_
15:**end for**
16:**return** *FeaSet*
~~~

### Code availability

After we reorganize the codes of EasiAI, the source codes will be available at github. Currently, it is available from on reasonable request.

### Data availability

The raw sEMG data supporting this study are not publicly available due to user privacy, but are available from the corresponding author on reasonable request. However, the data set, which are extracted from raw sEMG data by the feature extraction methods in this paper, support automatic CS identification study, are available at github after reorganized, each sample of which consist of 2949 features. Currently, it is available from on reasonable request.

## Acknowledgements

We thank Dr. Yunyou Huang for his help with the experimental design, data acquisition, data processing, and Fanda Fan for the help with the GBRT. We also thank Dr. Bingyan Cao and the other medical workers in Beijing Xiyuan Hospital rehabilitation physiotherapy for recruiting the subjects(the outpatients and the healthy free from the CS) and Research Center for Ubiquitous Computing Systems(in short CUbiCS) for the recruiting the subjects(the healthy). Besides, the authors also thank all subjects for providing valuable data. The paper was supported by the National Natural Science Foundation of China (NSFC) under Grant No. 61672498.

## Author contributions

N.-N.W designed the experiment and model, collected the data, wrote the codes, analyzed the data and wrote the manuscript. L.C. directed the project and revised the manuscript. X.H. guided the data preparation. Y.R. and J.X. participate in recruiting subjects and provide places for data acquisition. J.H.L participated in the data acquisition. N.W. provide the laptop and sEMG device.

## Correspondence

Correspondence and requests for materials should be addressed to Institute of Computing Technology, Chinese Academy of Sciences, No. 6 South Road of Academy of Sciences, Haidian District, Beijing 100089, China.

## Footnotes

1 The 421 features do not include FRR features.

2 The movements include *A*_1_, *A*_2_, *A*_3_, *A*_4_, *A*_5_, *A*_6_, *A*_7_.

3 The test set only includes the 282 features which are selected from the 2949 features by the EasiRF.

4 The feature number is the number of features extracted from muscles or movements. The more the features that are distributed on the muscle(movement) are, the stronger the muscle(movement) have the ability to identify CS, the more the muscle(movement) have differences between the healthy free from the CS and the CS suffer, the more the muscle(movement) is related to CS.

5 The importance are the contribution to the performance of task above in the training model. The more important the features from the muscle(movement) are, the stronger the muscle(movement) have the ability to identify CS, the more the muscle(movement) have differences between the healthy free from the CS and the CS suffer, the more the muscle(movement) is related to CS.

6 The imaging methods include spinal angiography, vertebral artery angiography, X-ray, computed to-mography(CT), and magnetic resonance imaging(MRI), etc.

